# Enhanced Episodic Memory Following Putative Inhibition of the Amygdala via Transcranial Low-Intensity Focused Ultrasound

**DOI:** 10.1101/2025.03.27.645800

**Authors:** Manoj K. Doss, Sydney R. Lambert, Jason Samaha, Charles B. Nemeroff, Gregory A. Fonzo, Joseph E. Dunsmoor

## Abstract

The amygdala is considered crucial to the formation of emotional episodic memories, but causal evidence in humans is limited due to challenges in non-invasive neuromodulation of deep brain structures. In a double-blind, sham-controlled, repeated measures study, we examined whether transcranial low-intensity focused ultrasound (tFUS) targeting the left amygdala prior to the encoding of emotional and neutral pictures impacted memory for these pictures 24 hours later. We used a putative inhibitory tFUS protocol shown to attenuate amygdala blood-oxygenation-level-dependent signal, thus testing the hypothesis that pre-encoding amygdala inhibition diminishes emotional memory. Surprisingly, active vs. sham sonication enhanced multiple measures of neutral and emotional memory across two memory tests. A secondary test of amygdala function found that active sonication enhanced fear recognition in faces.

Computational modeling further supported these results. These findings motivate a novel conceptualization of the amygdala’s role in emotional episodic memory. Rather than enhancing memory via amplification of salient stimuli, the amygdala may instead act as a filter that attenuates the maintenance of non-salient stimuli in long-term memory. Finally, the potential to enhance memory serves as an impetus to test tFUS of the amygdala in disorders such as depression and posttraumatic disorder that exhibit comorbid hyperreactive amygdalae and memory impairments.

## Introduction

The amygdala is crucial for myriad emotional processes, including the long-term retention of emotional episodic memories^1–4^. Episodic memory is the hippocampal-dependent conscious retrieval of information from the past, such as where or when one experienced an event. Emotional episodic memories, particularly emotionally negative memories, are typically remembered better than neutral memories (but see ^5^), and patients with amygdala lesions exhibit preferential memory impairments for emotional stimuli^6–10^. Furthermore, heightened amygdala activity has been found during the encoding^2^ (i.e., formation) and retrieval^3^ (i.e., remembering) of emotional memories. The amygdala is also thought to support the consolidation (i.e., post-encoding stabilization) of emotional episodic memories via interactions with the hippocampus and large-scale brain networks^4^. Consistent with the amygdala’s role in consolidation, one study in intracranial patients found that direct amygdala stimulation immediately after the presentation of neutral stimuli enhanced memory for these stimuli one day later but not during an immediate test^11^.

Although these lines of evidence implicate a critical role for the amygdala in emotional episodic memory, causal evidence in humans is largely lacking. Studies on patients with amygdala lesions cannot ascribe amygdala function to a specific phase of memory processing (i.e., encoding, consolidation, retrieval), and patients with lesions or those undergoing deep brain stimulation likely manifest major compensatory processes (cf. ^10^). While transcranial magnetic stimulation has become a common tool for non-invasively modulating brain activity, its effects cannot directly target deep brain structures^12^.

Transcranial low-intensity focused ultrasound (tFUS) is a relatively new technology that can reversibly and focally modulate neural activity by emitting converging high-frequency sound waves. Importantly, this approach is less hindered by energy degradation over distance, thereby allowing for the modulation of brain activity in deeper structures not easily accessible to non-invasive electromagnetic neuromodulation methods^13^. One tFUS protocol (see Methods) targeting the right amygdala during functional magnetic resonance imaging (fMRI) has been found to attenuate blood-oxygen-level-dependent (BOLD) amygdala activity and its functional connectivity with regions of the default mode network^14^, a critical network for episodic memory^15^. Likewise, this tFUS protocol targeting the left amygdala shortly before a fear-inducing task reduced amygdala and hippocampus reactivity, as well as functional connectivity between these regions^16^, suggesting that the effects of tFUS persist following online sonication. In terms of behavior, however, one study found that this tFUS protocol targeting the right amygdala increased subjective arousal ratings of emotionally negative pictures^17^ that were accompanied by increases in heart rate. To date, no studies have examined the effects of tFUS on human episodic memory.

Here, we tested the effects of a previously employed tFUS protocol demonstrating consistent evidence for amygdala inhibitory effects^14,16^. We sonicated the left amygdala immediately prior to the encoding phase of an emotional episodic memory task containing emotionally negative, neutral, and positive pictures. We chose to target the left amygdala, as some evidence suggests greater involvement of the left compared to right amygdala in emotional processing^18^. Memory was tested 24 hours post-sonication to mitigate potential carryover effects of sonication on retrieval. We hypothesized that the typical benefit in memory for emotionally negative pictures would be attenuated by down modulating the amygdala. Moreover, we expected such attenuations to be found in hippocampal-dependent recollection rather than cortical-dependent familiarity, as the amygdala is thought to support emotional recollection^2^.

Recollection is the retrieval of specific details (e.g., where or when an event took place), whereas familiarity is knowing that a stimulus has been processed without necessarily remembering corroborating evidence (e.g., recognizing a face without remembering how this individual is known)^19^. Finally, we administered a dynamic emotional facial expression task as a secondary measure to examine the impact of amygdala down modulation on emotional recognition. We hypothesized that down modulating the amygdala would impair recognition of fearful faces, given that amygdala lesions can diminish sensitivity to fear in faces^1^.

## Methods

### Participants

Thirty neurotypical young adults were recruited from The University of Texas at Austin and the surrounding area, but two participants did not complete the study following amygdala targeting, leaving 28 participants (10 males, age: *M* = 23.57, *SD* = 7.64). This *N* is greater than or equal to similar neuromodulation investigations of episodic memory^20^. Inclusion criteria were 18-50 years, and exclusion criteria were use of medications (other than birth control), current psychiatric, neurological, or developmental disorders, and MRI contraindications. Participants were asked to refrain from alcohol and other psychoactive drugs (other than caffeine) 24 hours prior to a testing day. All participants provided informed consent, and the study was approved by The University of Texas at Austin Institutional Review Board (IRB# STUDY00005037).

### Study Design

This study used a double-blind, sham-controlled, repeated measures design with two experimental arms. Prior to experimental sessions, participants completed a practice version of the memory task and an MRI-guided targeting session for individualized transducer placement optimized for line-of-sight with the left amygdala. During the first experimental session of each arm, pre-sonication affective and physiological measures were obtained, followed by the administration of sham or active sonication, post-sonication physiological measures, the encoding phase of the memory task, the dynamic emotional facial expression task, post-sonication affective measures, and blinding measures. Twenty-four hours later, participants returned for a second experimental session in which memory was tested for stimuli from the encoding phase. The first sessions from each arm were separated by ≥14 days, and sonication order was counterbalanced across participants (sonication order did not interact with the effects of sonication).

### Transcranial Low-Intensity Focused Ultrasound

The tFUS system used in this study was the Brainsonix Pulsar 1002^21^, and the sonication protocol used here (10 minutes in 30 s on/off blocks) has been found to inhibit the amygdala during online sonication^14^ and shortly afterward during threat processing^16^. Stimulation parameters included a 10 Hz pulse repetition frequency, 5% duty cycle, 5 ms pulse width, 14.4 W/cm^2^ derated spatial peak pulse average intensity, 719.91 mW/cm^2^ derated spatial peak temporal average intensity, .64 MPa derated instantaneous peak pressure, and 650 kHz fundamental frequency. The sham pad was indistinguishable but contained a porous air-filled foam layer through which ultrasound cannot be transmitted^22^.

### Affective, Physiological, and Blinding Measures

To determine whether tFUS evoked any noticeable effects, we measured subjective positive and negative affect using the Positive and Negative Affect Schedule^23^ (PANAS), heart rate, and blood pressure before and after sonication. To assess blinding, we asked participants at the end of the sonication session which sonication condition they thought they (active, sham) received and their confidence (5-point scale, 1 = definitely sham, 2 = possibly sham, 3 = don’t know, 4 = possibly active, 5 = definitely active).

### Emotional Episodic Memory Task

This task is similar to that used in previous investigations of neuromodulation and emotional episodic memory^24–27^ (Figure 1). Stimuli consisted of 360 images from the International Affective Picture System^28^ and the Nencki Affective Picture System^29^ and three- to five-word labels describing these images (e.g., “woman mourning mother’s death”). Negative, neutral, and positive images were selected to have valence ratings between 2-4, 4.5-5.5, and 6-8, respectively. These images were divided into 4 sets (A-D) each containing 90 images (30 of each emotion condition) with similar valence and arousal ratings for emotion conditions. Effort was made to balance the semantic content (e.g., people, animals, tools) between emotion conditions and stimulus sets, as semantic relatedness can influence memory^30,31^. Images were rotated through sonication and target/lure conditions (see below) across participants in a Latin square design. All phases lasted 15-20 minutes, contained 180 pseudorandomized trials (no more than 3 stimuli of a given emotion in a row), and 2000-3000-ms intertrial intervals.

**Figure 1.**
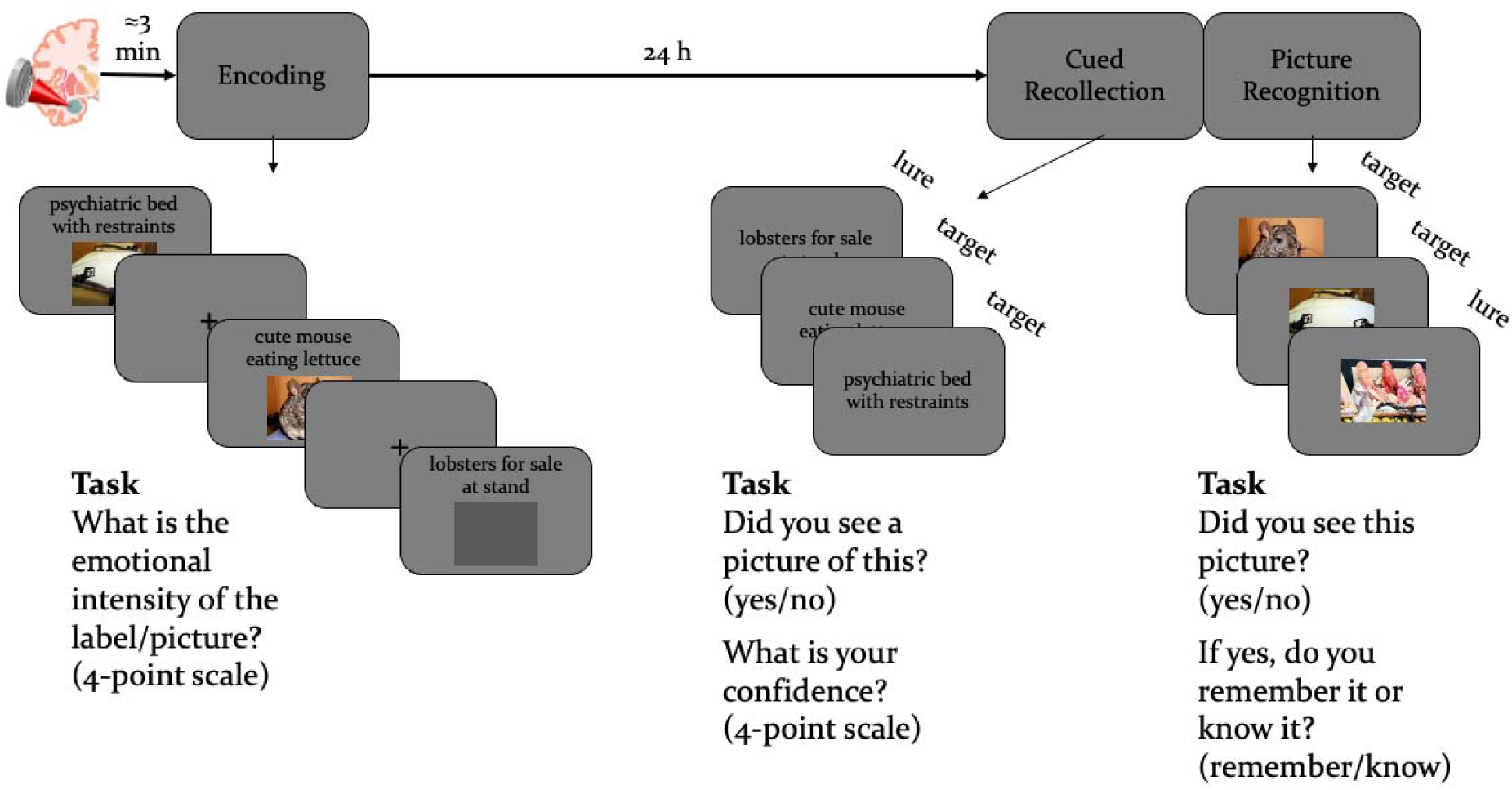
Emotional episodic memory task containing an encoding phase separated 24 hours later by two memory tests, a cued recollection test and picture recognition test. tFUS was administered shortly before the encoding phase.

On each trial of the encoding phase, participants were presented with a label for 3000 ms that was above its corresponding picture on half the trials or a grey box on the other half. Participants were to make arousal ratings (4-point scale) for labels and labels+pictures while they were on the screen. Twenty-four hours later, participants completed two self-paced memory tests, a cued recollection test and picture recognition test. On each trial of the cued recollection test, participants were presented with a label from the encoding phase for 1000 ms during which a response could not be made to allow enough time for recollection^19^. Afterward, participants responded whether a picture had been paired with this label (yes/no) followed by a 4-point confidence rating (1 = guessing, 4 = certain). Thus, targets were labels from encoding that had been presented with their corresponding pictures, and lures were labels from encoding that had been presented without their corresponding pictures during encoding.

Following the cued recollection test, participants completed the picture recognition test containing pictures from the encoding phase (targets) and pictures of the labels from the encoding phase presented without their corresponding picture (lures). On each trial, a picture was presented for 1000 ms before a response could be made. Afterward, participants responded whether the picture had been presented during the encoding phase (yes/no) followed by the remember/know procedure for estimating recollection and familiarity^32,33^. A “remember” response was to be made if specific details associated with the image could be recalled such as thoughts during its presentation, whereas a “know” response was to be made if the stimulus was known to presented in the absence of recalling any details.

### Dynamic Emotional Facial Expression Task

This task contained 108 randomized trials in which a video of a neutral face morphed over 1 second into different levels of fear or disgust (17%, 33%, 50%, 67%, 83%, 100%) or remained neutral (in which case the mouth simply opened; stimuli obtained from ^34^). In each emotion condition, there were 36 trials with 6 of each morph level for fear and disgust conditions and 6 identities equally distributed across these conditions. Participants were asked to determine whether the face was neutral, fear, or disgust. The same stimuli were used across experimental arms. This task was self-paced with 500-ms intertrial intervals and lasted 5-10 minutes.

### Data Analysis

Data from this study can be found at https://doi.org/10.17605/OSF.IO/YT8AX. For all analyses, α = .05 (two-tailed for pairwise contrasts), and significant or trending interactions were followed with paired sample *t* tests. Positive and negative PANAS scores, heart rate, and blood pressure were analyzed with 2 (sonication: sham, active) × 2 (timepoint: pre- vs. post-sonication) ANOVAs. One participant’s heart rate and blood pressure could not be collected. Blinding assessments for categorical responses and confidence were analyzed with separate χ^2^ tests (with a correction for continuity) for each sonication session and with 2 (sonication: sham, active) × 2 (sonication order: sham first, active first; sonication order was included, as participants may be better able to recognize the effects of tFUS after experiencing one condition), respectively.

Blinding measures were added to the study after data collection began, and thus, blinding data were collected for 14 and 16 participants in their first and second sonication sessions, respectively.

On the episodic memory task, 8 participants had <50% responses in at least one condition of the encoding phase, and 5 participants had negative memory accuracy in at least one condition of the cued recollection task, suggesting not following instructions. They were thus excluded, though inclusion of such noisy participants did not qualitatively change our results (Supplemental Material, SM). For the encoding phase, average arousal ratings were analyzed with 2 (sonication: sham, active) × 3 (emotion: negative, neutral, positive) × 2 (item: target, lure) repeated measures ANOVAs. Performance on the cued recollection and picture recognition tests was analyzed with 2 (sonication: sham, active) × 3 (emotion: negative, neutral, positive) repeated measures ANOVAs. For the cued recollection test, we calculated hit rates (*p*[“yes”|target]), false alarm rates (*p*[“yes”|lure]), accuracy (*p*[“yes”|target] - *p*[“yes”|lure]), and high-confidence versions of these measures by using only “yes” responses given the highest level of confidence. For the picture recognition test, we calculated hit rates, false alarm rates, and memory accuracy across “remember” and “know” responses and separately for “remember” and “know” responses. “Remember” responses are thought to reflect recollection, whereas “know” responses reflect familiarity in the absence of recollection when the two processes can co-occur. To account for unmeasured familiarity during “remember” responses for hit and false alarm rates, an independence remember/know (IRK) correction must be implemented by dividing *p*(“know”|target) and *p*(“know”|lure) by 1 - *p*(“remember”|target) and 1 - *p*(“remember”|lure), respectively^33,35,36^. We calculated IRK familiarity accuracy by subtracting IRK familiarity false alarm rates from IRK familiarity hit rates. To avoid dividing by 0, *p*(“remember”|target) and *p*(“remember”|lure) at floor and ceiling were replaced with .5/*N* and 1 - .5/*N*, respectively, where *N* is the number of trials^37^.

To estimate recollection and familiarity from the cued recollection test, confidence data were submitted to a dual-process signal detection (DPSD) analysis^19,38^ using the ROC toolbox for Matlab^39^. Briefly, hit rates are plotted against false alarm rates in a cumulative fashion starting with the highest level of confidence (i.e., the most conservative criterion) to the lowest level of confidence (i.e., the most lax criterion) with the final point always being (1,1). A receiver operator characteristic (ROC) curve is fit to these points using maximum likelihood estimation, but unlike typical ROC curves, the *y*-intercept is allowed to vary. The *y*-intercept of this function is thought to reflect the probability of accurate recollection, whereas the curvilinearity of the function is thought to reflect the strength of accurate familiarity. Due to a low trial count per condition/participant for model-fitting, which can produce unlikely results such as 0 estimates of recollection or familiarity, we performed a previously used bootstrapping procedure^20,24,36^. For each sonication × emotion condition, *N* participants were sampled with replacement 10,000 times, where *N* is the number of participants. For each sample, confidence counts were summed and fitted with the DPSD model to generate distributions of recollection and familiarity estimates. Contrasts between sonication conditions for each emotion were then made by subtracting two distributions to compute a confidence interval of this difference distribution and a *p*-value (proportion greater than 0).

One participant did not complete the dynamic emotional facial expression task, and 7 participants had <50% hit rates on 100% fear or disgust stimuli, suggesting not following instructions. They were thus excluded, though inclusion of such noisy participants did not qualitatively change our results (SM). Dependent variables included proportion correct (*p*[“fear”|fear) and *p*[“disgust”|disgust]) and incorrect (*p*[“fear”|disgust] and *p*[“disgust”|fear]). Correct identification was analyzed with 2 (sonication: sham, active) × 3 (emotion: fear, disgust) × 6 (morph: 17%, 33%, 50%, 67%, 83%, 100%) repeated measures ANOVAs. Incorrect identification was analyzed with 2 (sonication: sham, active) × 7 (morph: 0%, 17%, 33%, 50%, 67%, 83%, 100%, where 0% reflects neutral stimuli).

Due to the large number of comparisons across conditions and responses, “fear” and “disgust” responses were fit to separate four-parameter Naka–Rushton models using least squares estimation^40^:

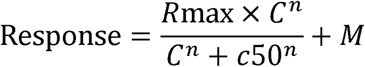

where *C* is the morph level (0%, 17%, 33%, 50%, 67%, 83%, 100%; 0% reflects neutral stimuli), *R*max is the saturation point (where the function plateaus), *c*50 is the threshold (the morph level at half the saturation point), *n* is the function’s slope, and *M* is an overall offset added to the response. A change in correct performance at the highest morph levels would be reflected in *R*max. A change in correct performance at moderate morph levels would be reflected in *c*50 or *n*, and a bias to respond “fear” or “disgust” would be reflected in *M*. Fitting this model without parameter constraints led to impossible parameters (e.g., *R*max > 1), and thus we constrained *R*max, *c*50, *n*, and *M* to .4-1, .5-10, .1-.8, and 0-.5, respectively. The same bootstrapping procedure and statistical contrasts conducted for DPSD modeling was performed on each parameter.

## Results

### Subjective Ratings and Physiological Data

Sonication did not modulate PANAS scores (positive: *F*(1, 27) = .62, *p* > .250; negative: *F*(1, 27) = .35, *p* > .250), heart rate (*F*(1, 26) = .52, *p* > .250), or blood pressure (systolic: *F*(1, 26) = .18, *p* > .250; diastolic: *F*(1, 26) = 2.10, *p* > .160). There were trending/significant main effects of timepoint for positive PANAS scores (*F*(1, 27) = 34.67, *p* < .001, 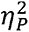= .56) and heart rate (*F*(1, 26) = 3.21, *p* = .085, 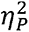 = .11), as both decreased within a session. Participants were at chance in determining whether they received active vs. sham sonication based on their categorical responses (first session: χ^2^(1) = 0, *p* > .250; second session: χ^2^(1) = 0, *p* > .250) and confidence (*F*(1, 12) = .03, *p* > .250). All other main effects and interactions were non-significant (all *F*s < 2.50, all *p*s > .100).

### Encoding Arousal Ratings

See SM for table of arousal ratings from the encoding phase. Negative stimuli were rated as more arousing than neutral and positive stimuli, and positive stimuli were rated as more arousing than neutral stimuli (*F*(2, 38) = 90.22, *p* < .001, 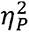= .83). Targets (labels+pictures) were rated as more arousing than lures (labels only; *F*(1, 19) = 8.21, *p* = .011, 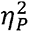= .29), and a trending emotion by item interaction (*F*(2, 38) = 2.85, *p* = .070, 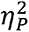= .13) suggested that this effect was more robust for negative and positive stimuli. All other main effects and interactions were non-significant (all *F*s < 2.50, all *p*s > .100).

### Cued Recollection Performance

Figure 2 displays performance on the cued recollection test (also see table in SM). Memory was best for negative pictures, as evidenced by main effects of emotion for hit rates (*F*(2, 44) = 25.70, *p* < .001, 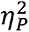= .54), accuracy (*F*(2, 44) = 27.84, *p* < .001, 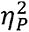= .56), high-confidence hit rates (*F*(2, 44) = 42.44, *p* < .001, 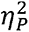= .66), and high-confidence accuracy (*F*(2, 44) = 39.37, *p* < .001, 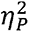= .64). High-confidence false alarm rates tended to be greatest for negative stimuli (*F*(2, 44) = 3.20, *p* = .051, 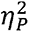= .13), though note that high-confidence false alarm rates were near floor. While the effect of sonication was non-significant for accuracy (*F*(1, 22) = 2.03, *p* = .168, 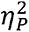= .08), surprisingly, active sonication at encoding compared to sham enhanced/tended to enhance hit rates (*F*(1, 22) = 3.11, *p* = .091, 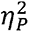= .12), high-confidence hit rates (*F*(1, 22) = 5.75, *p* = .025, 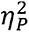= .21), and high-confidence accuracy (*F*(1, 22) = 5.96, *p* = .023, 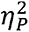= .21). These enhancements were most apparent for negative and neutral stimuli, though the sonication by emotion interactions were non-significant as were other main effects/interactions (all *F*s < 2.00, all *p*s > .150).

**Figure 2.**
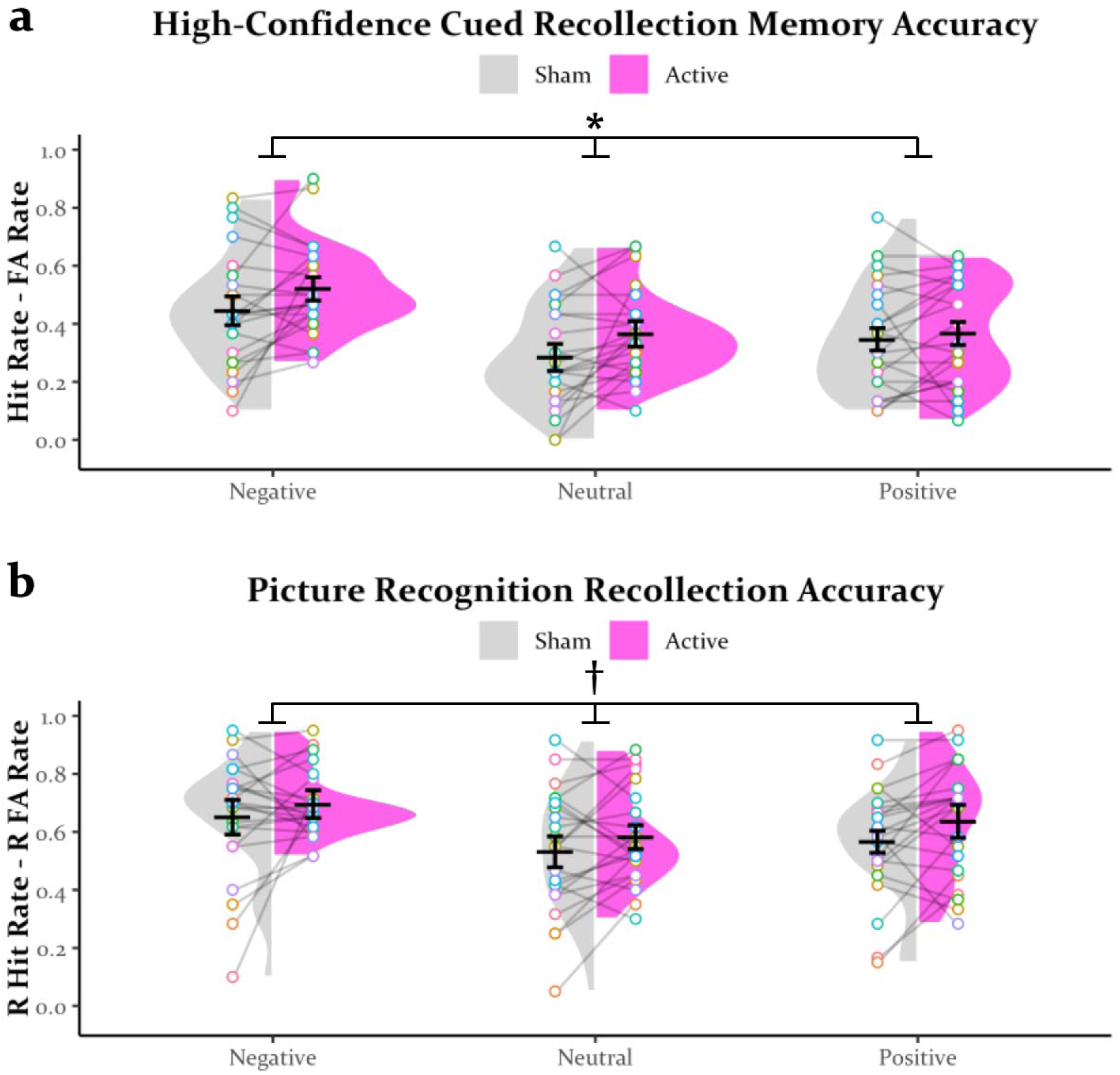
High-confidence accuracy (a) on the cued recollection test and recollection (i.e., remember) accuracy (b) on the picture recognition test. * = main effect of sonication (*p* < .050), † = trending main effect of sonication (*p* < .100), FA = false alarm, R = remember.

### DPSD Modeling

Figure 3 displays the aggregate ROC curves and distributions of recollection and familiarity estimates from DPSD modeling. Estimates of recollection were zero-inflated (a problem that can arise from high-confidence false alarms and less use of moderate confidence responses) and thus should be treated with caution. Active sonication did not significantly modulate recollection for negative (*M* =.06, *SD* = .10, *CI* = [-.14, .23], *p* >.250) or positive (*M* = .07, *SD* = .08, *CI* = [-.07, .24], *p* = .228) stimuli, though it enhanced recollection for neutral stimuli (*M* =.15, *SD* = .07, *CI* = [.00, .29], *p* = .026). Regarding familiarity, there were significant/trending enhancements of active sonication for negative (*M* = .25, *SD* = .16, *CI* = [-.04, .56], *p* = .051) and positive (*M* = .28, *SD* = .16, *CI* = [-.01, .60], *p* = .031) stimuli but not neutral stimuli (*M* = .11, *SD* = .14, *CI* = [-.15, .37], *p* = .201).

**Figure 3.**
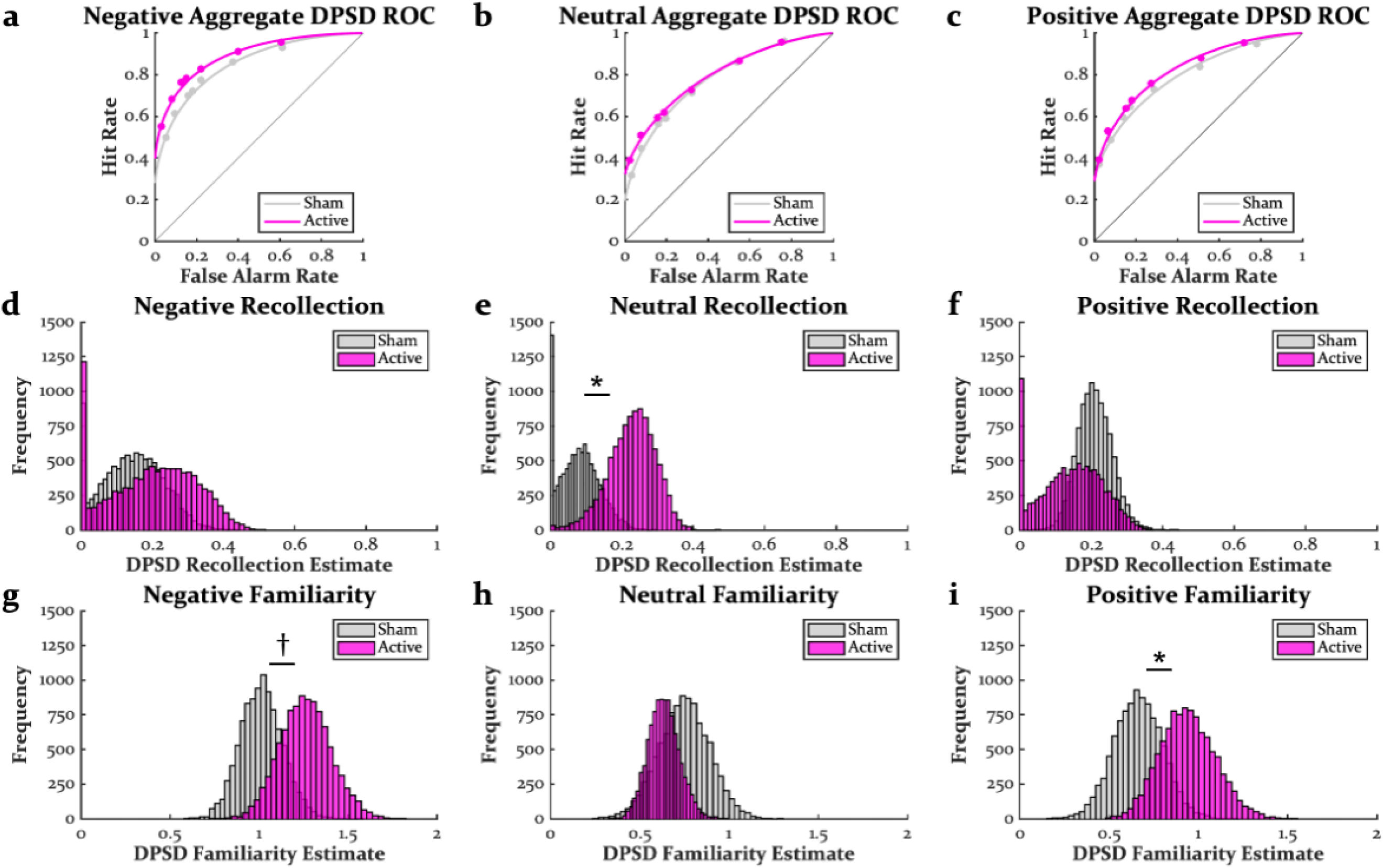
Aggregate receiver operator characteristic (ROC) curves (a-c), bootstrap distributions of recollection estimates (d-f), and bootstrap distributions of familiarity estimates (g-i) from dual process signal detection (DPSD) modeling. * = effect of sonication (*p* < .050), † = trending effect of sonication (p < .100).

### Picture Recognition Performance

Figure 2 displays performance on the picture recognition test (also see table in SM). Memory was best for negative pictures, as evidenced by main effects of emotion for hit rates (*F*(2, 44) = 8.48, *p* < .001, 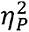= .28), accuracy (*F*(2, 44) = 11.24, *p* < .001, 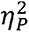= .34), remember hit rates (*F*(2, 44) = 13.02, *p* < .001, 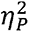 = .37), and remember accuracy (*F*(2, 44) = 12.57, *p* < .001, 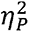 = .36). There was a trending effect of emotion for IRK familiarity false alarm rates (*F*(2, 44) = 2.76, *p* = .074, 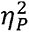= .11), which were greatest for neutral stimuli. Compared to sham, active sonication tended to enhance hit rates (*F*(1, 22) = 3.64, *p* = .069, 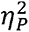= .14), remember hit rates (*F*(1, 22) = 3.74, *p* = .066, 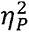= .15), and remember accuracy (*F*(1, 22) = 4.01, *p* = .058, 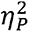= .15). There was a trending sonication by emotion interaction for hit rates (*F*(2, 44) = 2.88, *p* =.067, 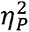= .12), as memory enhancements were most apparent for positive stimuli. All other main effects/interactions were non-significant (all *F*s < 2.50, all *p*s > .100).

### Dynamic Emotional Facial Expression Performance

Figure 4 displays performance on the dynamic emotional facial expression task (also see table in SM). Correct identification was better at higher morph levels (*F*(5, 95) = 136.49, *p* < .001, 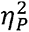= .88) and for disgust compared to fear (*F*(1, 19) = 28.29, *p* < .001, 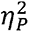= .60). Disgust was also more readily correctly identified at lower morph levels, as evidenced by an emotion by morph interaction (*F*(5, 95) = 8.05, *p* < .001, 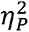= .30). There was a trending effect of sonication (*F*(1, 19) = 4.02, *p* = .059, 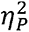= .17) and a significant sonication by morph interaction (*F*(5, 95) = 4.25, *p* = .002, 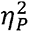= .18) on correct identification, as active sonication enhanced identification at moderate morph levels. Although the sonication by emotion (*F*(1, 19) = .60, *p* > .250) and three-way interaction (*F*(5, 95) = 1.38, *p* = .239) were non-significant for correct identification, Figure 4 highlights how this enhancement was most apparent for fearful faces. Incorrect “fear” and “disgust” responses were near floor, signifying that incorrect responses to fear and disgust stimuli were typically made with “neutral” responses. Incorrect “disgust” responses to fear stimuli tended to be greater than incorrect “fear” responses to disgust stimuli (*F*(1, 19) = 4.37, *p* = .050, 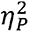= .19), and this trend was qualified by an emotion by morph interaction (*F*(6, 114) = 6.46, *p* < .001, 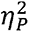 = .25), as more incorrect “disgust” responses were made at higher levels of fear. Although the effect of sonication was non-significant (*F*(1, 19) = .61, *p* > .250), there was a sonication by emotion interaction (*F*(1, 19) = 8.93, *p* = .008, 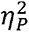 = .32), as incorrect “disgust” responses were somewhat greater under active sonication. All other main effects/interactions were non-significant (all *F*s < 1.50, all *p*s > .150).

**Figure 4.**
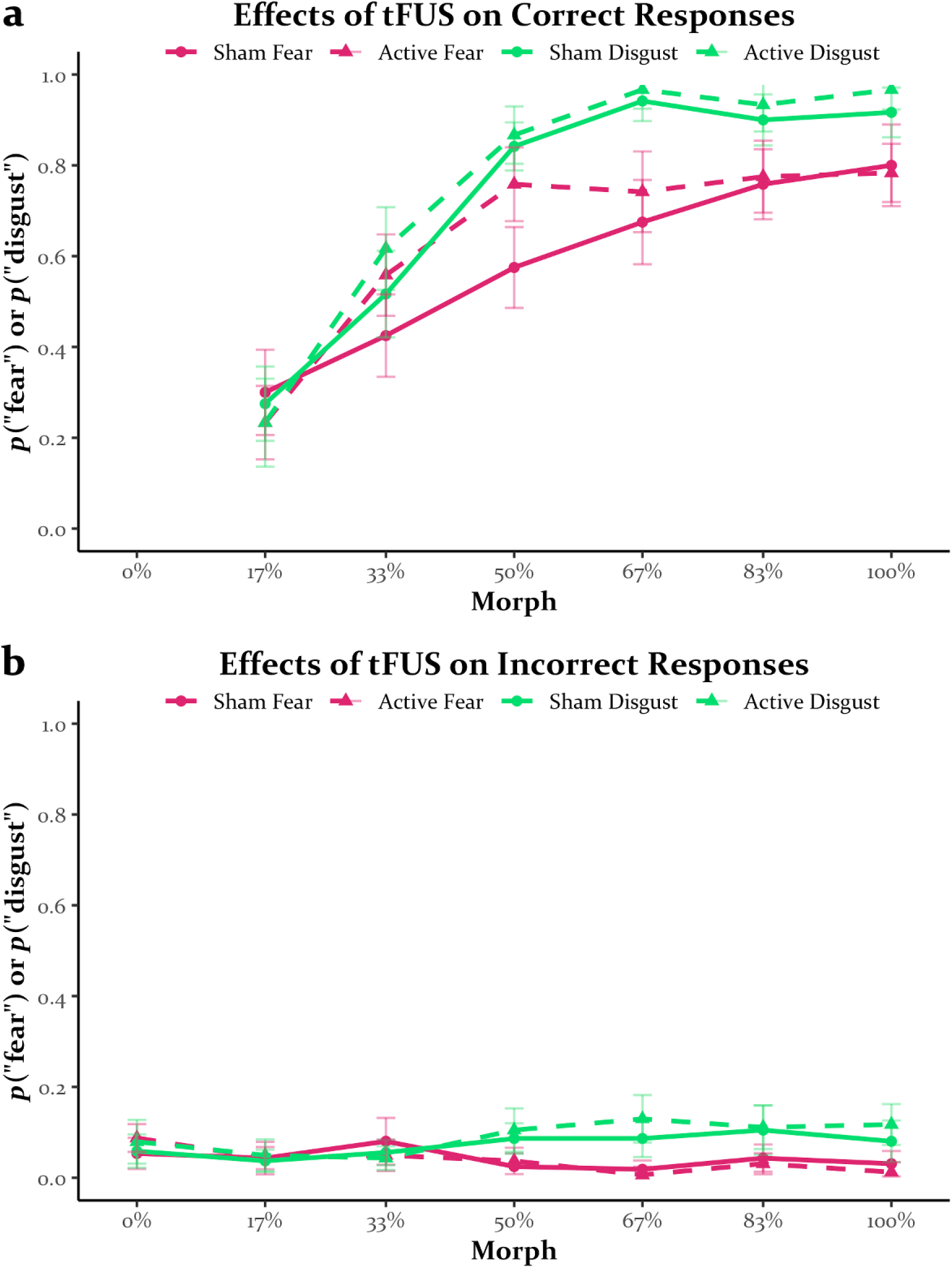
Correct (a) and incorrect (b) responses on the dynamical emotional facial expression task. Note that for incorrect responses, the color of the line denotes the response that was made, not the stimulus that was presented.

### Naka-Rushton Modeling

Figure 5 displays the aggregate Naka-Rushton fits and distributions of parameter estimates. The distribution of the saturation point for fearful faces in the sham condition was largely confined to 1 and thus could not be analyzed. For fearful faces, active sonication increased the slope (*M* = 2.23, *SD* = .72, *CI* = [1.22, 3.57], *p* < .001) and offset (*M* = .04, *SD* = .02, *CI* = [.00, .07], *p* = .026) while decreasing the threshold (*M* = .19, *SD* = .08, *CI* = [.01, .32], *p* = .017). Note that a decrease in threshold signifies improved detection at lower morph levels and a steeper slope indicates greater sensitivity to changes in fearful emotional content. For disgust faces, active sonication did not impact the saturation point (*M* = .03, SD = .05, *CI* = [-.06, .13], *p* > .250), threshold (*M* = .01, *SD* = .02, *CI* = [-.04, .06], *p* > .250), or offset (*M* = .03, *SD* = .03, *CI* = [-.02, .08], *p* = .114), but it tended to increase the slope (*M* = .73, *SD* = .60, *CI* = [-.27, 2.01], *p* =.064).

**Figure 5.**
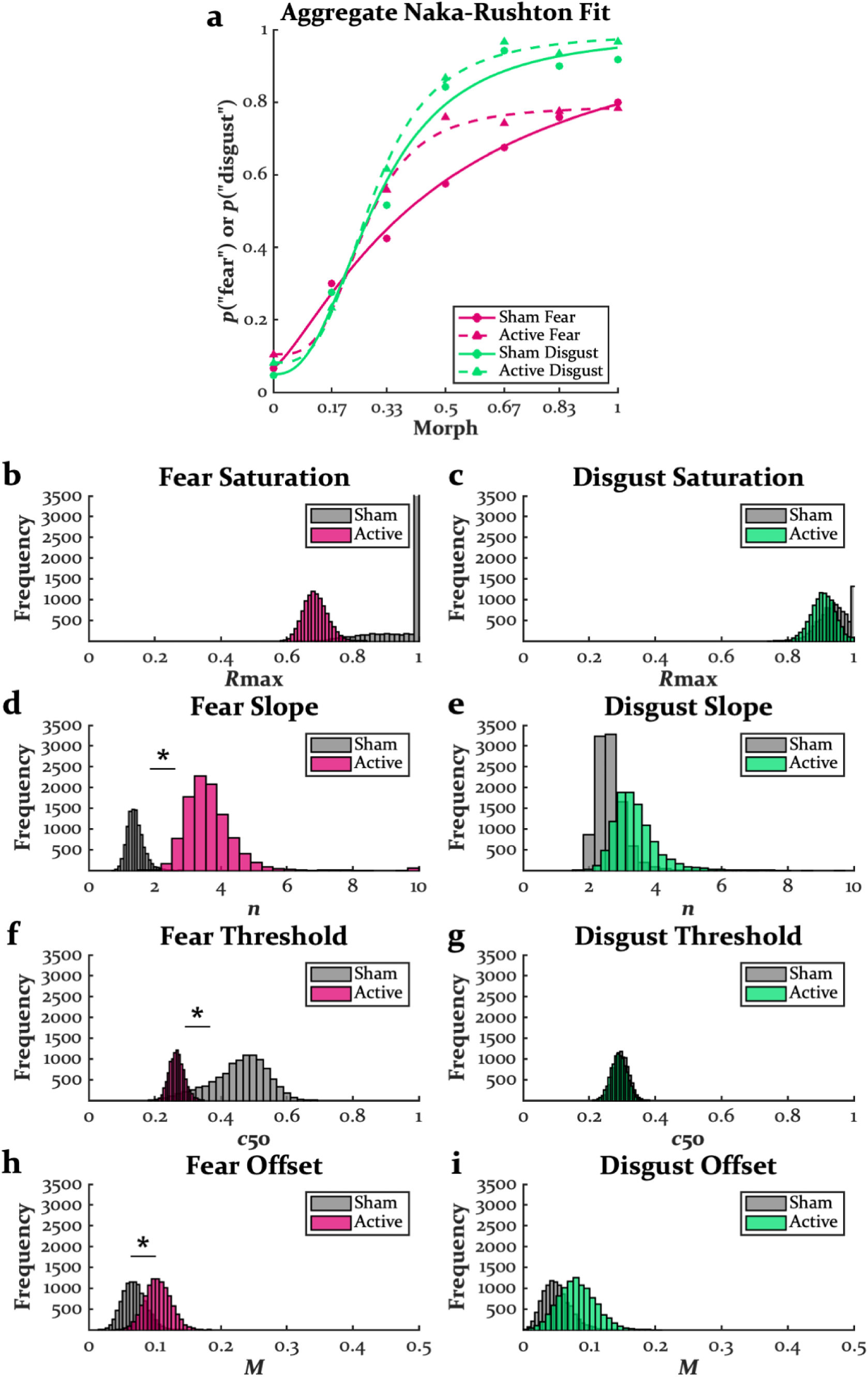
Aggregate Naka-Rushton fits (a), bootstrap distributions of the saturation point (b-c), threshold (d-e), slope (f-g), and offset (h-i). * = effect of sonication (*p* < .050).

## Discussion

The human amygdala has long been thought to directly mediate processes underlying the enhancement of emotional memory. This understanding is based on seminal findings from rodent models^41^, which has been translated using human neuroimaging research^42^. However, evidence that the human amygdala is necessary for the modulatory effects of emotion on memory has remained largely correlative. Here, we found that putative inhibitory tFUS of the left amygdala enhanced various metrics of both emotional and neutral memory, contrary to our initial hypothesis. These enhancements tended to come from hit rates rather than false alarm rates, suggesting that a greater number of stimuli were remembered rather than changes in memory monitoring processes^43^. Moreover, these enhancements were found in both hippocampal-dependent recollection and cortical-dependent familiarity^19^. This is the first demonstration of the effects of tFUS in the human brain on episodic memory. We also found that putative inhibitory tFUS of the amygdala enhanced emotional recognition with preferential effects for fearful compared to disgust faces. This finding was also unexpected given that amygdala lesions can impair fear recognition^1^, though such effects may be less reliable with unilateral lesions^44^ and lesions in adulthood^45^. This enhancement of fear recognition was not due to an overall bias, as misattribution of fear was low. In contrast, lesions and stimulation of the amygdala have been found to lead to misattributions of emotion to facial stimuli^7,46^.

One explanation for these unexpected effects is that the protocol we used did not in fact inhibit the amygdala but instead facilitated its activity. For example, although this protocol has been found to inhibit the amygdala during online sonication^14^, there could be a rebound effect immediately following sonication (i.e., when participants completed tasks). However, one study that used this same sonication protocol found that BOLD signal in the amygdala remained less active during a fear inducing task shortly after sonication^16^. Alternatively, by inhibiting only the left amygdala, the right amygdala could have overcompensated, though these other tFUS studies did not report increases in contralateral BOLD activity in the amygdala. Other regions may have overcompensated such as the insula, which is highly interconnected with the amygdala and also plays a critical role in emotion recognition^47,48^. While another study also found surprising effects of left amygdala sonication using a similar tFUS protocol, namely increased subjective arousal ratings to negative pictures and increased heart rate^17^, we did not observe such effects.

Another explanation for these findings is that instead of the amygdala enhancing the processing of emotional stimuli, it may instead filter out the processing of less salient stimuli. Although both these accounts would predict that memory should be relatively better for emotional stimuli compared to neutral stimuli, the latter would suggest that by attenuating the amygdala’s filtering function, there should be memory enhancements for both neutral and emotional stimuli, as even some emotional stimuli could be filtered. Attenuation of such filtering should also result in greater sensitivity to emotional facial expressions when less emotional information is provided. Indeed, the amygdala has been suggested to act as a sort of filter^49^, with evidence coming from memory impairments for neutral stimuli that immediately precede negative stimuli and neutral stimuli that immediately proceed negative or positive stimuli^50–52^.

Such filtering of neutral memories in the context of emotional stimuli is abolished by amygdala lesions^53^. Moreover, a recent study in patients with intracranial electrodes found that excitatory stimulation of the amygdala immediately after a neutral stimulus was presented enhanced memory for these stimuli. A filtering account would suggest that heightening the amygdala’s filtering function immediately following the presentation of a stimulus could reduce retroactive interference, thereby allowing pre-stimulation memories to better stabilize. This explanation is similar to the retrograde facilitation effect observed following post-encoding administration of GABA_A_ positive allosteric modulators like alcohol^36,54^.

A limitation to neuromodulation investigations of memory is that the effects typically persist following encoding, and thus, could impact consolidation (cf. ^54,55^). Moreover, one possibility is that amygdala inhibition grew larger (for unbeknownst reasons) after encoding. If the amygdala does indeed bias encoding toward salient stimuli, a delayed inhibition effect would reduce retroactive interference. fMRI during encoding and post-encoding and inhibition of the amygdala immediately post-encoding could disentangle these different accounts. Finally, it could also be that inhibition of the amygdala persisted 24 hours later during retrieval. Although it is unclear why inhibition of the amygdala during retrieval should enhance memory when the amygdala increases its activation during emotional memories^3^, a possible explanation could be that amygdala inhibition homogenized psychological states between encoding and retrieval (i.e., state-dependent memory). Nevertheless, there was no evidence that tFUS had any subjective effects.

Many psychiatric disorders, including depression and posttraumatic stress disorder, exhibit a hyperreactive amygdala^56–58^ and impairments in episodic memory^59,60^, consistent with an ostensible filtering function of the amygdala. One study in two patients found that unilateral ablation of the amygdala to treat epilepsy simultaneously treated comorbid PTSD symptoms and improved episodic memory^46^. Recently, we found that 15 sessions of left amygdala tFUS over 3 weeks with the same protocol used here alleviated negative affect symptoms in patients that included those with depression and PTSD (unpublished). Thus, a potential secondary effect could be enhancing memory in these populations that exhibit comorbid memory impairments.

Such memory enhancements could even synergize with psychotherapy by improving retention of what is learned, providing an impetus to test inhibitory tFUS of the amygdala on episodic memory in clinical populations.

## Author Contributions

M.K.D. and J.E.D. conceptualized the study. M.K.D. designed the task, analyzed the data, and wrote the initial draft of the manuscript. S.R.L. collected the data. G.A.F. and C.B.N. provided support in administering tFUS. J.E.D. funded the study. M.K.D., S.R.L., G.A.F., and C.B.N. edited the manuscript.

## Acknowledgements

M.K.D. receives research support from the Effie and Wofford Cain Foundation and Center for MINDS. J.E.D. received funding for this project from the Health Transformation Research Institute (HTRI) at The University of Texas at Austin Dell Medical School. J.E.D. receives research support from the National Institutes of Health (NIH), National Science Foundation (NSF), and US-Israel Binational Science Foundation. G.A.F. receives research support from the NIH, Defense Advanced Research Projects Agency (DARPA), SEAL Future Foundation, Effie and Wofford Cain Foundation, and Center for MINDS. C.B.N. receives research support from NIH and Texas Child Mental Health Care Consortium. These funding sources (other than that from HTRI) had no role in this study other than supporting salaries.

