## Supplemental Materials for "Enhanced Episodic Memory Following Putative Inhibition of the Amygdala via Transcranial Low-Intensity Focused Ultrasound"

**Methods**

*Transducer Placement*

During the targeting session, participants underwent MRI at The University of Texas at Austin Biomedical Imaging Center (BIC) on a 3 T Siemens MAGNETOM Vida scanner using a 20-channel head coil. A T1-weighted high-resolution anatomical image (3D MPRAGE, sagittal, TR/TE = 2400/2.18 ms, slice thickness = .8 mm, flip angle = 8°, FOV = 256 mm, matrix size = 200 × 308 × 320, voxel size = .8 × .8 × .8 mm, duration = 6:38) was first acquired for use with the ANT Neuro visor2 Neuronavigation system (eemagine GmbH, Berlin, Germany). Personalized head models were rendered in the visor2 software using each participant’s T1 anatomical. During this time, the participant was removed from the scanner and escorted to the MR control room to have the tFUS transducer affixed to the left temporal window using an initial placement approximately halfway between the orbital socket and the ear with the middle of the transducer approximately parallel with the center of the eye. The T1-weighted image was used to provide an initial approximation regarding whether a 55 or 65 mm focal-depth transducer would be best for targeting the left amygdala. The transducer was held in place on the skull using a 3D-printed transducer holder and three Velcro straps that were attached around the forehead, chin, and between the forehead and chins straps. An ultrasound-transmitting transducer pad was placed between the transducer face and the side of the participant’s skull with a thin layer of ultrasound transmission gel between each contact point. Hair around the location of the transducer face was sprayed down with water and flattened to minimize the possibility of air bubbles between the transducer pad and scalp.

Following this placement of the transducer, participants were placed back inside the scanner. The head was tilted slightly to the right as needed to allow for the transducer and participant to fit comfortably within the head coil. A T1-weighted anatomical scout image (3D Fast Low Angle Shot [FLASH] with non-selective excitation, sagittal, TR/TE = 3.20/1.37 ms, slice thickness = 1.6 mm, flip angle = 8°, FOV = 260 mm, matrix size = 128 × 160 × 160, voxel size = 1.6 × 1.6 × 1.6 mm, duration = 0:17) was then collected to visualize the location of the transducer relative to the left amygdala. The center of the transducer has a circular ring that appears with bright intensity on T1-weighted sequences. The scanner image navigator crosshairs were centered within this ring in the sagittal anatomical scout image, and the axes of the crosshairs were then visually shifted by hand so that the angle of the crosshairs was parallel with the face of the transducer in the sagittal plane and perpendicular with the face of the transducer in the axial and coronal planes. This crosshair perpendicular to the transducer face approximates the ultrasound beam trajectory through skull and into the brain. By visual inspection, it was then determined if the transducer placement would provide a “line of sight” to the left amygdala. If the placement was off, the scanner software measurement tool was used to approximate the distance between the current line of sight from the center of the transducer to the left amygdala by drawing a line perpendicular to the crosshair-approximated line of sight to the center of the left amygdala in the axial and coronal planes. The participant was then removed from the scanner, and the placement of the transducer was adjusted via soft tape measure and surgical marker to match the offsets noted from the prior anatomical scout. The participant then returned to the scanner, and another anatomical scout was acquired to examine the new adjusted placement. This process was repeated until the line of sight from the transducer center was visually verified to lie within the center of the left amygdala. Visual verification of adequate transducer placement for line-of-sight-based targeting was verified by both the primary and secondary scanner operators, who were trained by GAF and JED to visually distinguish the left amygdala as the circular almond-shaped structure in the left medial temporal lobe (anterior to the hippocampus in the axial plane, separated by a thin strip of darker intensity voxels, lateral to the higher-intensity midbrain and cerebral peduncle, and just posterior to the piriform/olfactory cortex and posterior orbitofrontal cortex).

Once the transducer placement was established, participants were removed from the scanner and seated ≈1 m from the ANT visor 2 Neuronavigation system in the MRI operator suite. A 3D-printed attachment was used to attach the ANT visor2 optical tracker to the outside center of the transducer, which was calibrated a priori in the ANT visor2 system software using a custom focused ultrasound transducer software setting that allows for the specification of focal depth. Each participant’s head was then calibrated in the system software by attaching the head tracker to an adhesive lead pad on the bridge of the nose, and the typical method of using an optical pointer to locate nasion, inion, and the tragus of each ear was employed. Head shape points were then traced on the participants scalp while avoiding the Velcro straps, and the individualized head model based upon the participant’s MPRAGE was used to visualize the location of the transducer in relation to the amygdala target and programmed focal depth. If there was a large discrepancy between the transducer’s line-of-site target in the neuronavigation software and that verified via MR placement, the calibration process was repeated until there was visual agreement. A custom-made button-box was used to send a TTL trigger to the neuronavigation software, thereby marking the location of the transducer on each participant’s head model for repeated placement in experimental sessions.

**Results with Participant Exclusion**

**Table S1**

|  | Sham | Active |
| --- | --- | --- |
| *Encoding* |  |  |
| Target Neg | 2.87 ± .12, [2.61, 3.12] | 2.78 ± .11, [2.54, 3.02] |
| Target Neu | 1.63 ± .08, [1.46, 1.80] | 1.66 ± .08, [1.48, 1.83] |
| Target Pos | 2.14 ± .11, [1.91, 2.37] | 2.18 ± .11, [1.94, 2.42] |
| Lure Neg | 2.72 ± .11, [2.49, 2.94] | 2.48 ± .11, [2.26, 2.71] |
| Lure Neu | 1.55 ± .07, [1.41, 1.69] | 1.54 ± .07, [1.40, 1.67] |
| Lure Pos | 1.91 ± .10, [1.70, 2.13] | 1.95 ± .12, [1.70, 2.19] |
| *Cued Recollection* |  |  |
| Hit Rate Neg | .72 ± .03, [.65, .79] | .78 ± .03, [.73, .84] |
| Hit Rate Neu | .59 ± .04, [.51, .67] | .62 ± .03, [.56, .68] |
| Hit Rate Pos | .65 ± .04, [.57, .73] | .68 ± .04, [.60, .76] |
| FA Rate Neg | .18 ± .03, [.12, .24] | .15 ± .03, [.10, .20] |
| FA Rate Neu | .20 ± .03, [.14, .25] | .19 ± .03, [.13, .25] |
| FA Rate Pos | .17 ± .02, [.12, .22] | .18 ± .03, [.12, .24] |
| Accuracy Neg | .54 ± .04, [.45, .63] | .63 ± .03, [.56, .70] |
| Accuracy Neu | .39 ± .04, [.30, .48] | .43 ± .03, [.36, .50] |
| Accuracy Pos | .47 ± .04, [.39, .56] | .50 ± .05, [.40, .60] |
| Hi Hit Rate Neg | .50 ± .04, [.41, .59] | .55 ± .03, [.48, .62] |
| Hi Hit Rate Neu | .32 ± .04, [.24, .39] | .39 ± .03, [.32, .46] |
| Hi Hit Rate Pos | .37 ± .04, [.29, .45] | .39 ± .04, [.31, .47] |
| Hi FA Rate Neg | .05 ± .01, [.03, .08] | .03 ± .01, [.01, .05] |
| Hi FA Rate Neu | .03 ± .01, [.00, .06] | .02 ± .01, [.01, .04] |
| Hi FA Rate Pos | .02 ± .01, [.01, .04] | .02 ± .01, [.01, .04] |
| Hi Accuracy Neg | .44 ± .04, [.36, .53] | .52 ± .03, [.45, .59] |
| Hi Accuracy Neu | .28 ± .04, [.21, .36] | .37 ± .03, [.30, .43] |
| Hi Accuracy Pos | .35 ± .04, [.26, .43] | .37 ± .04, [.29, .45] |
| *Recognition* |  |  |
| Hit Rate Neg | .87 ± .03, [.81, .93] | .89 ± .02, [.86, .93] |
| Hit Rate Neu | .80 ± .03, [.73, .88] | .86 ± .02, [.82, .90] |
| Hit Rate Pos | .82 ± .03, [.75, .89] | .90 ± .01, [.87, .93] |
| FA Rate Neg | .08 ± .02, [.05, .12] | .08 ± .02, [.05, .11] |
| FA Rate Neu | .09 ± .02, [.05, .13] | .11 ± .02, [.07, .14] |
| FA Rate Pos | .09 ± .02, [.05, .12] | .09 ± .02, [.05, .13] |
| Accuracy Neg | .79 ± .04, [.71, .87] | .81 ± .02, [.77, .86] |
| Accuracy Neu | .71 ± .04, [.64, .79] | .75 ± .03, [.69, .81] |
| Accuracy Pos | .73 ± .04, [.65, .81] | .80 ± .03, [.75, .86] |
| R Hit Rate Neg | .68 ± .04, [.60, .76] | .72 ± .02, [.67, .77] |
| R Hit Rate Neu | .56 ± .04, [.46, .65] | .61 ± .03, [.54, .68] |
| R Hit Rate Pos | .59 ± .04, [.51, .67] | .66 ± .04, [.58, .74] |
| R FA Rate Neg | .03 ± .01, [.02, .04] | .02 ± .00, [.02, .03] |
| R FA Rate Neu | .02 ± .00, [.02, .03] | .02 ± .00, [.02, .03] |
| R FA Rate Pos | .02 ± .00, [.02, .03] | .02 ± .00, [.02, .03] |
| R Accuracy Neg | .65 ± .04, [.56, .74] | .70 ± .02, [.65, .75] |
| R Accuracy Neu | .53 ± .04, [.44, .62] | .58 ± .03, [.51, .65] |
| R Accuracy Pos | .57 ± .04, [.48, .65] | .64 ± .04, [.55, .72] |
| K Hit Rate Neg | .19 ± .03, [.14, .25] | .17 ± .02, [.13, .22] |
| K Hit Rate Neu | .25 ± .03, [.18, .31] | .25 ± .03, [.20, .30] |
| K Hit Rate Pos | .23 ± .03, [.17, .29] | .24 ± .03, [.17, .31] |
| K FA Rate Neg | .07 ± .01, [.05, .10] | .07 ± .01, [.04, .10] |
| K FA Rate Neu | .09 ± .02, [.05, .12] | .10 ± .01, [.07, .13] |
| K FA Rate Pos | .08 ± .01, [.05, .11] | .09 ± .02, [.06, .13] |
| K Accuracy Neg | .12 ± .03, [.07, .17] | .10 ± .02, [.05, .15] |
| K Accuracy Neu | .16 ± .03, [.10, .23] | .15 ± .02, [.11, .20] |
| K Accuracy Pos | .15 ± .03, [.09, .21] | .15 ± .03, [.09, .21] |
| IRK F Hit Rate Neg | .65 ± .05, [.55, .75] | .62 ± .05, [.53, .72] |
| IRK F Hit Rate Neu | .57 ± .05, [.46, .68] | .64 ± .04, [.56, .72] |
| IRK F Hit Rate Pos | .57 ± .05, [.47, .67] | .68 ± .04, [.60, .76] |
| IRK F FA Rate Neg | .07 ± .01, [.05, .10] | .07 ± .01, [.04, .10] |
| IRK F FA Rate Neu | .09 ± .02, [.05, .12] | .10 ± .02, [.07, .13] |
| IRK F FA Rate Pos | .08 ± .01, [.05, .11] | .09 ± .02, [.06, .13] |
| IRK F Accuracy Neg | .58 ± .05, [.48, .68] | .55 ± .05, [.45, .65] |
| IRK F Accuracy Neu | .48 ± .05, [.37, .59] | .54 ± .04, [.46, .62] |
| IRK F Accuracy Pos | .49 ± .05, [.38, .59] | .59 ± .04, [.50, .67] |

Performance on the emotional episodic memory task with 8 participants excluded from the encoding data for <50% responses and 6 participants excluded from the cued recollection and recognition data for negative memory accuracy on the cued recollection test. Values are mean ± standard error of the mean [95% confidence interval]. Neg = negative, Neu = neutral, Pos = positive, FA = false alarm, Hi = high-confidence, R = remember, K = know, IRK F = independence remember/know familiarity.

**Table S2**

|  | Sham | Active |
| --- | --- | --- |
| Correct "Fear" Fear 17% | .30 ± .05, [.19, .41] | .23 ± .04, [.16, .31] |
| Correct "Fear" Fear 33% | .42 ± .06, [.30, .55] | .56 ± .05, [.45, .66] |
| Correct "Fear" Fear 50% | .58 ± .05, [.46, .69] | .76 ± .04, [.67, .84] |
| Correct "Fear" Fear 67% | .68 ± .06, [.56, .79] | .74 ± .05, [.65, .84] |
| Correct "Fear" Fear 83% | .76 ± .04, [.68, .84] | .77 ± .04, [.68, .87] |
| Correct "Fear" Fear 100% | .80 ± .04, [.73, .87] | .78 ± .03, [.72, .85] |
| Correct "Disg" Disg 17% | .28 ± .04, [.19, .36] | .23 ± .05, [.13, .33] |
| Correct "Disg" Disg 33% | .52 ± .05, [.42, .62] | .62 ± .05, [.51, .73] |
| Correct "Disg" Disg 50% | .84 ± .03, [.77, .91] | .87 ± .04, [.79, .95] |
| Correct "Disg" Disg 67% | .94 ± .02, [.89, .99] | .97 ± .02, [.93, 1.00] |
| Correct "Disg" Disg 83% | .90 ± .03, [.84, .96] | .93 ± .03, [.88, .99] |
| Correct "Disg" Disg 100% | .92 ± .03, [.86, .98] | .97 ± .02, [.93, 1.00] |
| Incorrect "Fear" Neutral | .07 ± .02, [.03, .11] | .10 ± .02, [.06, .15] |
| Incorrect "Fear" Disg 17% | .05 ± .02, [.00, .10] | .03 ± .01, [.00, .05] |
| Incorrect "Fear" Disg 33% | .10 ± .03, [.03, .17] | .06 ± .02, [.01, .10] |
| Incorrect "Fear" Disg 50% | .03 ± .02, [.00, .07] | .04 ± .02, [.00, .08] |
| Incorrect "Fear" Disg 67% | .02 ± .01, [-.01, .04] | .01 ± .01, [-.01, .03] |
| Incorrect "Fear" Disg 83% | .04 ± .02, [.00, .08] | .03 ± .02, [-.01, .07] |
| Incorrect "Fear" Disg 100% | .03 ± .02, [.00, .07] | .02 ± .01, [-.01, .04] |
| Incorrect "Disg" Neutral | .05 ± .02, [.01, .09] | .08 ± .03, [.02, .14] |
| Incorrect "Disg" Fear 17% | .03 ± .01, [.00, .05] | .04 ± .02, [.00, .08] |
| Incorrect "Disg" Fear 33% | .04 ± .02, [.01, .08] | .03 ± .02, [.00, .07] |
| Incorrect "Disg" Fear 50% | .08 ± .02, [.04, .13] | .08 ± .02, [.04, .11] |
| Incorrect "Disg" Fear 67% | .06 ± .02, [.01, .11] | .12 ± .03, [.05, .18] |
| Incorrect "Disg" Fear 83% | .09 ± .03, [.03, .15] | .09 ± .03, [.03, .15] |
| Incorrect "Disg" Fear 100% | .05 ± .02, [.01, .09] | .10 ± .03, [.05, .15] |

Performance on the dynamic emotional facial expression task with 7 participants excluded for <50% hit rates on 100% fear and disgust stimuli. Values are mean ± standard error of the mean [95% confidence interval]. Disg = disgust.

**Results Without Participant Exclusion**

*Encoding Arousal Ratings*

Table S3 displays arousal ratings on the encoding phase of the emotional episodic memory task. Two participants had no responses in at least one condition, and thus, could not be analyzed. Negative stimuli were rated as more arousing than neutral and positive stimuli, and positive stimuli were rated as more arousing than neutral stimuli (*F*(2, 50) = 131.82, *p* < .001, $\eta_{p}^{2}$ = .84). Targets (labels+pictures) were rated as more arousing than lures (labels only; *F*(1, 25) = 12.57, *p* = .002, $\eta_{p}^{2}$ = .33), and a marginally trending emotion by item interaction (*F*(2, 50) = 12.57, *p* = .100, $\eta_{p}^{2}$ = .09) suggested that this effect was more robust for negative and positive stimuli. Although the main effect of sonication was non-significant (*F*(1, 25) = .29, *p* > .250), there was a trending sonication by emotion interaction (*F*(2, 50) = 2.52, *p* = .090, $\eta_{p}^{2}$ = .09), as arousal ratings for negative lures were attenuated by active sonication (95% CI: [.02, .31], *t*(25) = 2.32, *p* = .029, *d* = .45) but not in negative and positive conditions (all *t*s < 1.00, all *p*s > .250). Note, however, that such an effect was not found for negative targets (*t*(25) = .97, *p* > .250), but this effect on lures must be treated with caution given the non-significant three-way interaction (*F*(2, 50) = 1.24, *p* > .250). All other interactions were non-significant (all *F*s < 2.50, all *p*s > .200).

**Table S3**

|  | Sham | Active |
| --- | --- | --- |
| *Encoding* |  |  |
| Target Neg | 2.82 ± .10, [2.63, 3.02] | 2.75 ± .09, [2.56, 2.94] |
| Target Neu | 1.65 ± .06, [1.52, 1.78] | 1.67 ± .07, [1.53, 1.82] |
| Target Pos | 2.15 ± .09, [1.97, 2.33] | 2.20 ± .09, [2.02, 2.39] |
| Lure Neg | 2.67 ± .08, [2.50, 2.85] | 2.51 ± .09, [2.33, 2.68] |
| Lure Neu | 1.53 ± .06, [1.42, 1.65] | 1.54 ± .06, [1.43, 1.66] |
| Lure Pos | 1.92 ± .09, [1.74, 2.10] | 1.95 ± .10, [1.75, 2.14] |
| *Cued Recollection* |  |  |
| Hit Rate Neg | .73 ± .03, [.67, .79] | .77 ± .02, [.73, .82] |
| Hit Rate Neu | .60 ± .03, [.53, .67] | .62 ± .02, [.57, .67] |
| Hit Rate Pos | .64 ± .03, [.57, .71] | .68 ± .03, [.62, .75] |
| FA Rate Neg | .27 ± .04, [.18, .36] | .23 ± .05, [.14, .32] |
| FA Rate Neu | .27 ± .04, [.19, .36] | .26 ± .04, [.18, .35] |
| FA Rate Pos | .26 ± .04, [.17, .35] | .26 ± .04, [.17, .35] |
| Accuracy Neg | .46 ± .05, [.36, .56] | .54 ± .05, [.44, .65] |
| Accuracy Neu | .32 ± .05, [.22, .43] | .36 ± .04, [.27, .45] |
| Accuracy Pos | .38 ± .05, [.28, .49] | .43 ± .05, [.32, .53] |
| Hi Hit Rate Neg | .47 ± .04, [.39, .56] | .52 ± .04, [.44, .59] |
| Hi Hit Rate Neu | .30 ± .03, [.23, .37] | .35 ± .03, [.29, .42] |
| Hi Hit Rate Pos | .34 ± .04, [.27, .42] | .36 ± .04, [.29, .44] |
| Hi FA Rate Neg | .09 ± .02, [.04, .13] | .06 ± .02, [.01, .10] |
| Hi FA Rate Neu | .06 ± .02, [.02, .09] | .05 ± .02, [.01, .09] |
| Hi FA Rate Pos | .04 ± .01, [.02, .07] | .05 ± .02, [.01, .09] |
| Hi Accuracy Neg | .38 ± .04, [.30, .47] | .46 ± .04, [.37, .54] |
| Hi Accuracy Neu | .24 ± .04, [.16, .32] | .30 ± .04, [.23, .38] |
| Hi Accuracy Pos | .30 ± .04, [.22, .38] | .31 ± .04, [.23, .39] |
| *Recognition* |  |  |
| Hit Rate Neg | .85 ± .03, [.79, .91] | .88 ± .02, [.85, .92] |
| Hit Rate Neu | .80 ± .03, [.74, .87] | .85 ± .02, [.81, .89] |
| Hit Rate Pos | .81 ± .03, [.74, .87] | .89 ± .01, [.86, .92] |
| FA Rate Neg | .10 ± .02, [.06, .15] | .13 ± .03, [.06, .20] |
| FA Rate Neu | .11 ± .02, [.06, .16] | .14 ± .03, [.08, .20] |
| FA Rate Pos | .12 ± .03, [.07, .17] | .12 ± .02, [.07, .17] |
| Accuracy Neg | .75 ± .04, [.65, .84] | .75 ± .04, [.67, .84] |
| Accuracy Neu | .69 ± .04, [.61, .77] | .71 ± .04, [.64, .78] |
| Accuracy Pos | .69 ± .04, [.60, .78] | .77 ± .03, [.71, .83] |
| R Hit Rate Neg | .66 ± .04, [.57, .74] | .71 ± .02, [.66, .76] |
| R Hit Rate Neu | .55 ± .04, [.47, .63] | .58 ± .03, [.52, .65] |
| R Hit Rate Pos | .58 ± .04, [.50, .65] | .63 ± .03, [.56, .70] |
| R FA Rate Neg | .04 ± .01, [.01, .07] | .04 ± .01, [.02, .07] |
| R FA Rate Neu | .04 ± .01, [.01, .06] | .04 ± .01, [.01, .07] |
| R FA Rate Pos | .04 ± .01, [.01, .07] | .04 ± .01, [.02, .05] |
| R Accuracy Neg | .62 ± .05, [.52, .71] | .67 ± .03, [.60, .73] |
| R Accuracy Neu | .51 ± .04, [.43, .60] | .54 ± .04, [.47, .62] |
| R Accuracy Pos | .54 ± .04, [.45, .62] | .60 ± .04, [.52, .68] |
| K Hit Rate Neg | .19 ± .02, [.15, .24] | .17 ± .02, [.13, .21] |
| K Hit Rate Neu | .25 ± .03, [.20, .31] | .27 ± .03, [.21, .32] |
| K Hit Rate Pos | .23 ± .03, [.18, .29] | .26 ± .03, [.20, .32] |
| K FA Rate Neg | .08 ± .01, [.06, .10] | .10 ± .02, [.06, .14] |
| K FA Rate Neu | .09 ± .01, [.06, .12] | .11 ± .02, [.07, .15] |
| K FA Rate Pos | .09 ± .02, [.06, .13] | .10 ± .02, [.07, .13] |
| K Accuracy Neg | .11 ± .02, [.07, .16] | .07 ± .02, [.02, .12] |
| K Accuracy Neu | .16 ± .03, [.10, .21] | .15 ± .02, [.12, .19] |
| K Accuracy Pos | .14 ± .03, [.08, .19] | .16 ± .03, [.11, .21] |
| IRK F Hit Rate Neg | .62 ± .04, [.53, .71] | .61 ± .04, [.52, .69] |
| IRK F Hit Rate Neu | .58 ± .04, [.48, .67] | .63 ± .03, [.56, .70] |
| IRK F Hit Rate Pos | .57 ± .04, [.48, .66] | .69 ± .03, [.62, .76] |
| IRK F FA Rate Neg | .09 ± .01, [.06, .11] | .11 ± .03, [.06, .17] |
| IRK F FA Rate Neu | .10 ± .02, [.07, .13] | .12 ± .02, [.08, .17] |
| IRK F FA Rate Pos | .10 ± .02, [.06, .15] | .11 ± .02, [.07, .14] |
| IRK F Accuracy Neg | .53 ± .05, [.44, .63] | .49 ± .05, [.39, .59] |
| IRK F Accuracy Neu | .48 ± .05, [.38, .57] | .51 ± .04, [.44, .58] |
| IRK F Accuracy Pos | .46 ± .05, [.36, .56] | .58 ± .03, [.51, .65] |

Performance on the emotional episodic memory task without excluding participants. Values are mean ± standard error of the mean [95% confidence interval]. Neg = negative, Neu = neutral, Pos = positive, FA = false alarm, Hi = high-confidence, R = remember, K = know, IRK F = independence remember/know familiarity.

*Cued Recollection Performance*

Figure S1 and Table S3 display performance on the cued recollection test. Memory was best for negative pictures, as evidenced by main effects of emotion for hit rates (*F*(2, 54) = 25.48, *p* < .001, $\eta_{p}^{2}$ = .49), accuracy (*F*(2, 54) = 23.01, *p* < .001, $\eta_{p}^{2}$ = .46), high-confidence hit rates (*F*(2, 54) = 47.93, *p* < .001, $\eta_{p}^{2}$ = .43), and high-confidence accuracy (*F*(2, 54) = 36.18, *p* < .001, $\eta_{p}^{2}$ = .57). High-confidence false alarm rates were greatest for negative stimuli (*F*(2, 54) = 4.59, *p* = .014, $\eta_{p}^{2}$ = .15), though this effect should be treated with caution, as high-confidence false alarm rates were near floor. Compared to sham, active sonication at encoding surprisingly tended to enhance hit rates (*F*(1, 27) = 3.19, *p* = .085, $\eta_{p}^{2}$ = .11), high-confidence hit rates (*F*(1, 27) = 4.75, *p* = .038, $\eta_{p}^{2}$ = .15), and high-confidence accuracy (*F*(1, 27) = 3.95, *p* = .057, $\eta_{p}^{2}$ = .13), though this effect was non-significant for accuracy (*F*(1, 27) = 2.54, *p* = .123, $\eta_{p}^{2}$ = .09). These enhancements were most apparent for negative and neutral stimuli, though the sonication by emotion interactions were non-significant (hit rates: *F*(2, 54) = .41, *p* > .250, high-confidence hit rates: *F*(2, 54) = .71, *p* > .250; high-confidence accuracy: *F*(2, 54) = 2.27, *p* = .114, $\eta_{p}^{2}$ = .08). Further evidence of memory enhancement from sonication came from a sonication by emotion interaction for high-confidence false alarm rates (*F*(2, 54) = 3.85, *p* = .027, $\eta_{p}^{2}$ = .12), as sonication tended to decrease high-confidence false alarm rates to negative stimuli (95% CI: [.00, 07], *t*(27) = 1.80, *p* = .084, *d* = .34) but not neutral (95% CI: [-.03, .05], *t*(27) = .51, *p* > .250) or positive stimuli (95% CI: [-.02, .03], *t*(27) = .61, *p* > .250). However, note again that high-confidence false alarm rates were near floor. All other main effects/interactions were non-significant (all *F*s < 1.00, all *p*s > .250).

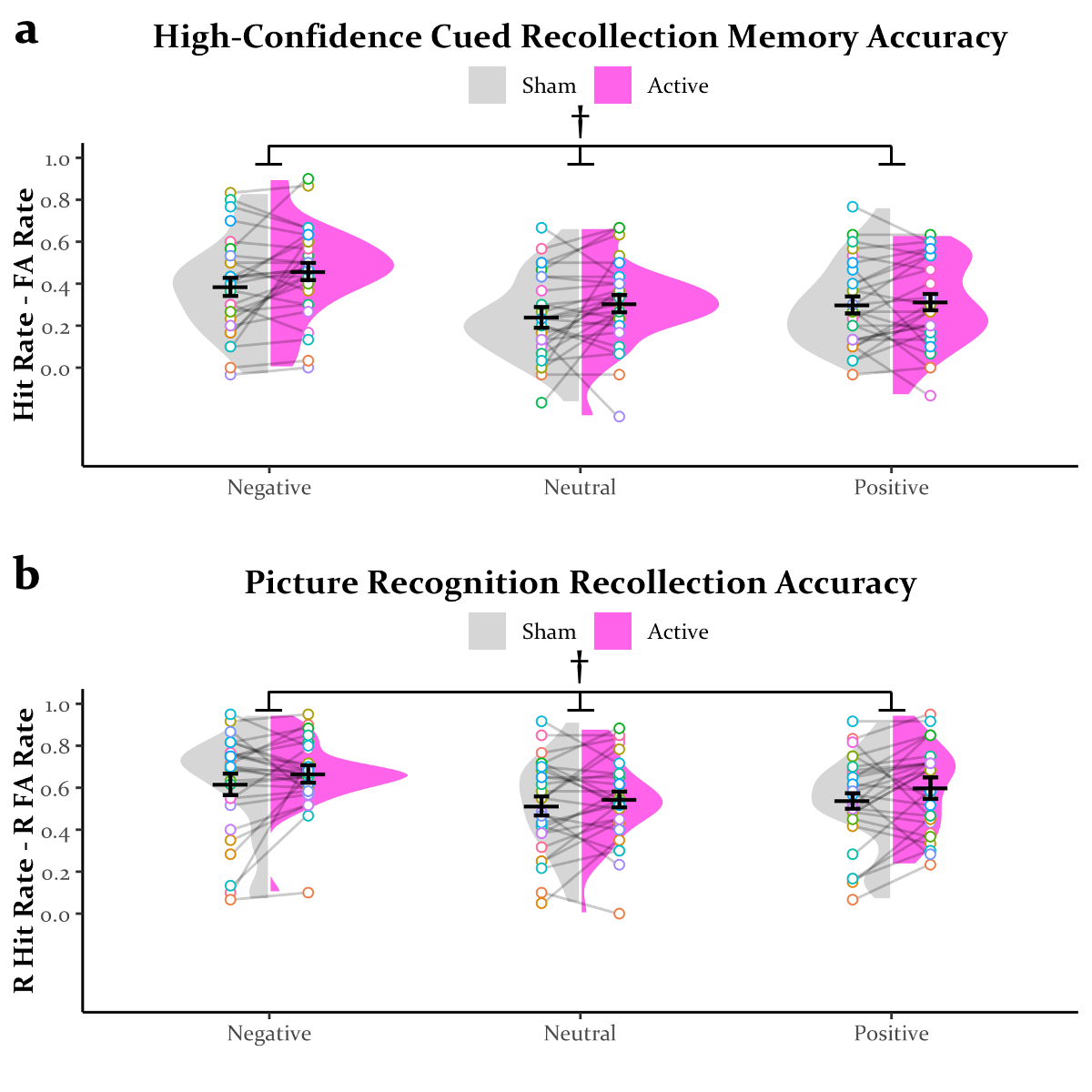

**Figure S1.** High-confidence accuracy (**a**) on the cued recollection test and recollection (i.e., remember) accuracy (**b**) on the picture recognition test. * = main effect of sonication (*p* < .050), † = trending main effect of sonication (*p* < .100), FA = false alarm, R = remember.

*DPSD Modeling*

Figure S2 displays the aggregate ROC curves and distributions of recollection and familiarity from DPSD modeling. Estimates of recollection were zero-inflated (a problem that can arise from high-confidence false alarms and less use of moderate confidence responses) and thus should be treated with caution. Active sonication did not significantly modulate recollection for negative (*M* = .10, *SD* = .09, 95% *CI* = [-.09, .27], *p* = .154) or positive (*M* = .09, *SD* = .08, *CI* = [-.04, .25], *p* = .125) stimuli, though there was a trend toward enhancement of recollection for neutral stimuli (*M* = .11, *SD* = .07, *CI* = [-.04, .25], *p* = .072). Active sonication did not significantly modulate familiarity for negative (*M* = .17, *SD* = .15, *CI* = [-.13, .46], *p* = .132) or neutral (*M* =.05, *SD* = .13, *CI* = [-.19, .30], *p* > .250) stimuli, though it enhanced familiarity for positive stimuli (*M* = .32, *SD* =.14, *CI* = [.06, .60], *p* = .006).

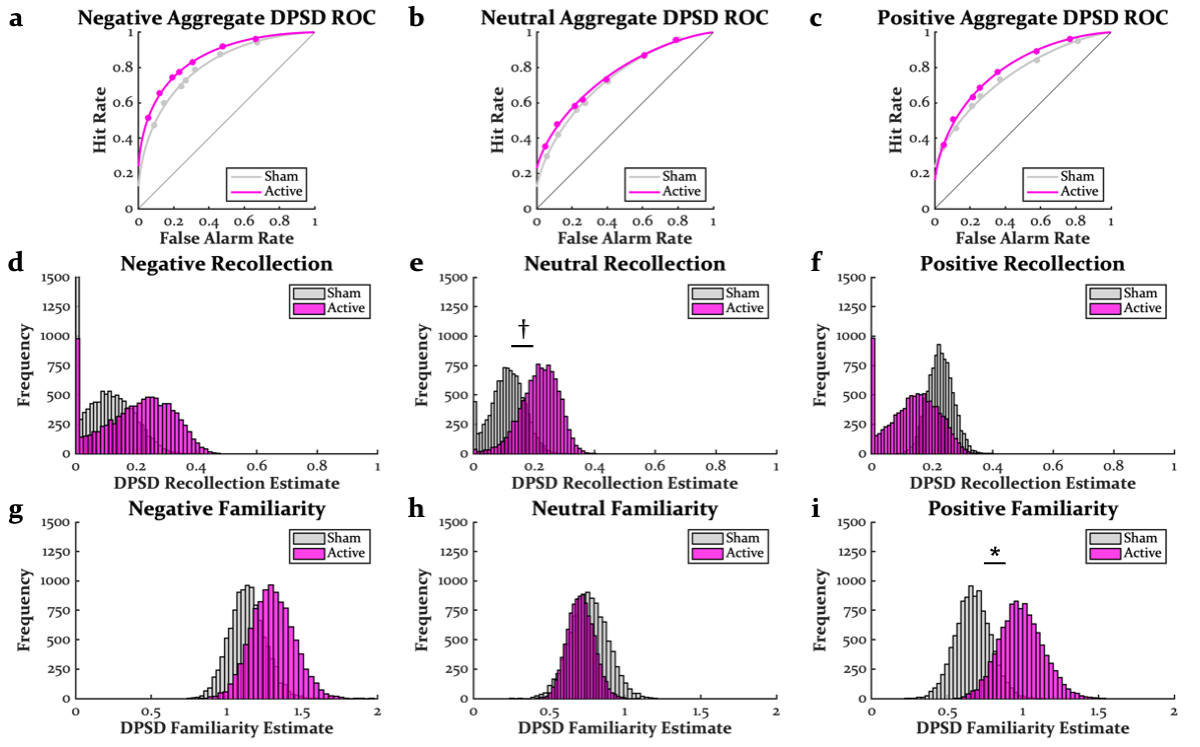

**Figure S2.** Aggregate receiver operator characteristic (ROC) curves (**a**-**c**), bootstrap distributions of recollection estimates (**d**-**f**), and bootstrap distributions of familiarity estimates (**g**-**i**) from dual process signal detection (DPSD) modeling. * = effect of sonication (*p* < .050), † = trending effect of sonication (*p* < .100).

*Picture Recognition Performance*

Figure S1 and Table S3 display performance on the picture recognition test. Memory was best for negative pictures, as evidenced by significant main effects of valence for hit rates (*F*(2, 54) = 5.58, *p* = .006, $\eta_{P}^{2}$ = .17), accuracy (*F*(2, 54) = 5.55, *p* = .006, $\eta_{P}^{2}$ = .17), remember hit rates (*F*(2, 54) = 16.50, *p* < .001, $\eta_{P}^{2}$ = .38), and remember accuracy (*F*(2, 54) = 15.95, *p* < .001, $\eta_{P}^{2}$ = .37). Compared to sham, active sonication tended to enhance hit rates (*F*(1, 27) = 5.59, *p* = .026, $\eta_{P}^{2}$ = .17), remember hit rates (*F*(1, 27) = 3.87, *p* = .060, $\eta_{P}^{2}$ = .05), remember accuracy (*F*(1, 27) = 3.56, *p* = .070, $\eta_{P}^{2}$ = .05), and IRK familiarity hit rates (*F*(1, 27) = 4.18, *p* = .051, $\eta_{P}^{2}$ = .13). Although the effects of sonication on accuracy (*F*(1, 27) = 1.48, *p* = .234) and IRK familiarity accuracy (*F*(1, 27) = 1.49, *p* = .233) were not significant, the sonication by emotion interaction was significant/trending for accuracy (*F*(2, 54) = 3.19, *p* = .049, $\eta_{P}^{2}$ = .11) and IRK familiarity accuracy (*F*(2, 54) = 2.96, *p* = .061, $\eta_{P}^{2}$ = .10), as well as for hit rates (*F*(2, 54) = 3.05, *p* = .056, $\eta_{P}^{2}$ = .10). These interactions were explained by significant/trending memory enhancements for positive stimuli (hit rates: 95% CI: [.03, .14], *t*(27) = 3.02, *p* = .005, *d* = .57; accuracy: 95% CI: [.01, .15], *t*(27) = 2.38, *p* = .025, *d* = .45; IRK familiarity accuracy: 95% CI: [.02, .22], *t*(27) = 2.43, *p* = .022, *d* = .46) and sometimes for neutral stimuli (hit rates: 95% CI: [.00, .10], *t*(27) = 1.94, *p* = .062, *d* = .37; accuracy: 95% CI: [-.05, .08], *t*(27) = .59, *p* > .250; IRK familiarity accuracy: 95% CI: [-.06, .13], *t*(27) = .80, *p* > .250), but not for negative stimuli (hit rates: 95% CI: [-.02, .09], *t*(27) = 1.28, *p* = .21; accuracy: 95% CI: [-.07, .09], *t*(27) = .19, *p* > .250; IRK familiarity accuracy: 95% CI: [-.07, .15], *t*(27) = .73, *p* > .250). All other main effects/interactions were non-significant (all *F*s < 1.50, all *p*s > .250).

*Dynamic Emotional Facial Expression Performance*

Figure S3 and Table S4 display performance on the dynamic emotional facial expression task. Correct identification was better at higher morph levels (*F*(5, 130) = 126.19, *p* < .001, $\eta_{P}^{2}$ = .83) and for disgust compared to fear (*F*(1, 26) = 38.79, *p* < .001, $\eta_{P}^{2}$ = .60). Disgust was also more readily correctly identified at lower morph levels, as evidenced by an emotion by morph interaction (*F*(5, 130) = 13.47, *p* < .001, $\eta_{P}^{2}$ = .34). There was a trending effect of sonication (*F*(1, 26) = 3.18, *p* = .086, $\eta_{P}^{2}$ = .11) and a significant sonication by morph interaction (*F*(5, 130) = 2.38, *p* = .042, $\eta_{P}^{2}$ = .08) on correct identification, as active sonication enhanced identification at moderate morph levels. Although the sonication by emotion (*F*(1, 26) = .25, *p* > .250) and three-way interaction (*F*(1, 26) = .76, *p* > .250) were non-significant for correct identification, Figure S3 highlights how this enhancement was most apparent for fearful faces. Incorrect “fear” and “disgust” responses were near floor, signifying that incorrect responses to fear and disgust stimuli were typically made with “neutral” responses. Incorrect “disgust” responses to fear stimuli were greater than incorrect “fear” responses to disgust stimuli (*F*(1, 26) = 9.72, *p* = .004, $\eta_{P}^{2}$ = .27), and this effect was qualified by an emotion by morph interaction (*F*(6, 156) = 8.68, *p* < .001, $\eta_{P}^{2}$ = .25), as more incorrect “disgust” responses were made at higher levels of fear. Although the effect of sonication was non-significant (*F*(1, 26) = .73, *p* > .250), there was a trending sonication by emotion interaction (*F*(1, 26) = 3.64, *p* = .068, $\eta_{P}^{2}$ = .12), as incorrect “disgust” responses were somewhat greater under active sonication. All other main effects/interactions were non-significant (all *F*s < 1.50, all *p*s > .250).

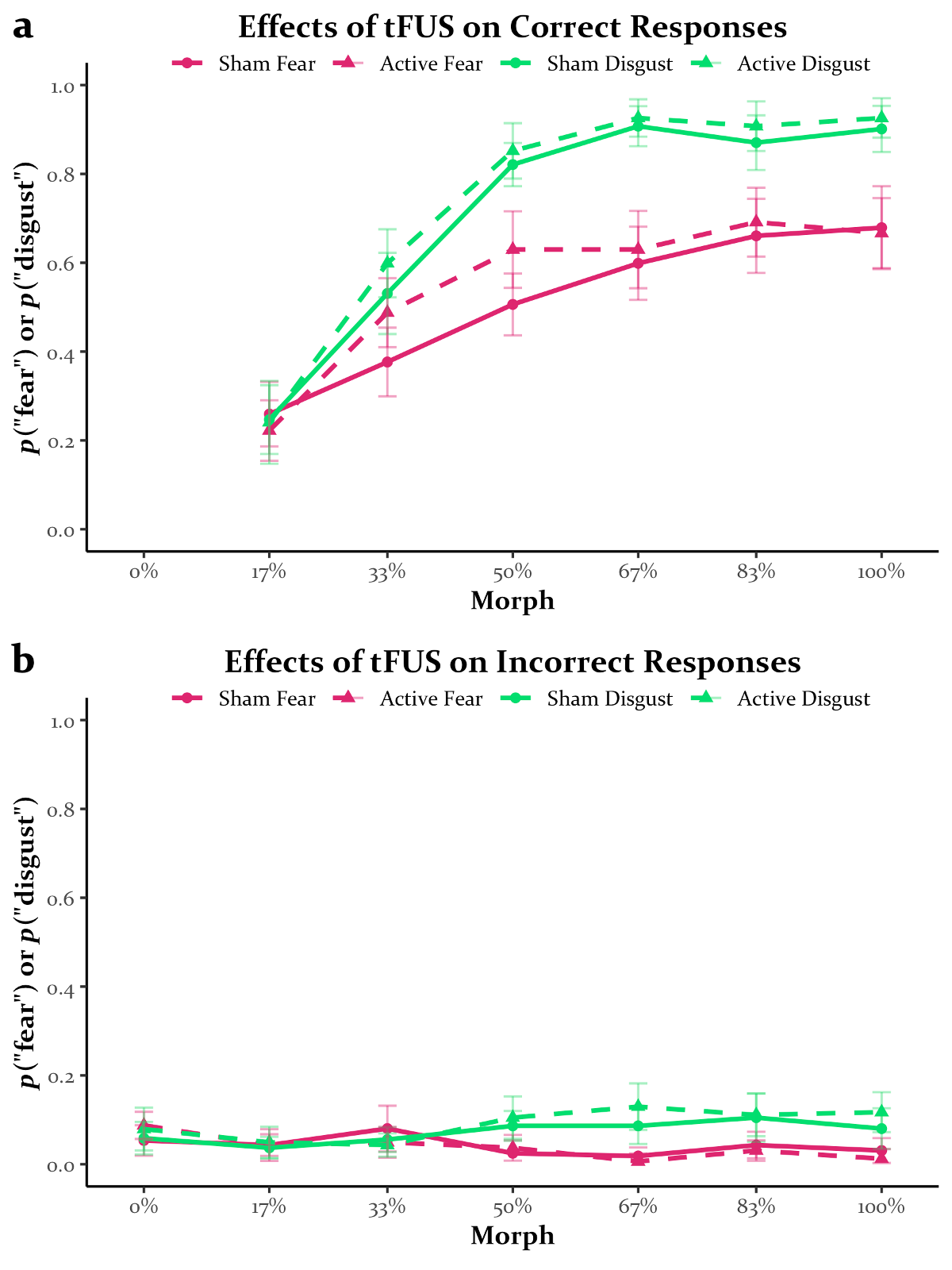

**Figure S3.** Correct (**a**) and incorrect (**b**) responses on the dynamical emotional facial expression task. Note that for incorrect responses, the color of the line denotes the response that was made, not the stimulus that was presented.

**Table S4**

|  | Sham | Active |
| --- | --- | --- |
| Correct "Fear" Fear 17% | .26 ± .04, [.17, .35] | .22 ± .03, [.16, .29] |
| Correct "Fear" Fear 33% | .38 ± .05, [.27, .48] | .49 ± .05, [.39, .59] |
| Correct "Fear" Fear 50% | .51 ± .05, [.40, .61] | .63 ± .06, [.51, .74] |
| Correct "Fear" Fear 67% | .60 ± .06, [.48, .71] | .63 ± .06, [.51, .74] |
| Correct "Fear" Fear 83% | .66 ± .05, [.55, .77] | .69 ± .05, [.58, .80] |
| Correct "Fear" Fear 100% | .68 ± .06, [.56, .79] | .67 ± .05, [.56, .77] |
| Correct "Disg" Disg 17% | .25 ± .03, [.18, .32] | .24 ± .04, [.16, .33] |
| Correct "Disg" Disg 33% | .53 ± .04, [.45, .61] | .60 ± .05, [.50, .70] |
| Correct "Disg" Disg 50% | .82 ± .03, [.76, .89] | .85 ± .04, [.78, .93] |
| Correct "Disg" Disg 67% | .91 ± .03, [.84, .98] | .93 ± .03, [.87, .98] |
| Correct "Disg" Disg 83% | .87 ± .04, [.79, .95] | .91 ± .03, [.84, .97] |
| Correct "Disg" Disg 100% | .90 ± .03, [.84, .96] | .93 ± .03, [.86, .99] |
| Incorrect "Fear" Neutral | .05 ± .01, [.02, .08] | .09 ± .02, [.05, .12] |
| Incorrect "Fear" Disg 17% | .04 ± .02, [.00, .08] | .04 ± .02, [.01, .08] |
| Incorrect "Fear" Disg 33% | .08 ± .03, [.03, .13] | .05 ± .02, [.01, .09] |
| Incorrect "Fear" Disg 50% | .02 ± .01, [.00, .05] | .04 ± .02, [.00, .07] |
| Incorrect "Fear" Disg 67% | .02 ± .01, [.00, .04] | .01 ± .01, [-.01, .02] |
| Incorrect "Fear" Disg 83% | .04 ± .02, [.01, .08] | .03 ± .02, [.00, .06] |
| Incorrect "Fear" Disg 100% | .03 ± .01, [.00, .06] | .01 ± .01, [-.01, .03] |
| Incorrect "Disg" Neutral | .06 ± .02, [.02, .10] | .08 ± .03, [.03, .13] |
| Incorrect "Disg" Fear 17% | .04 ± .02, [.00, .07] | .05 ± .02, [.01, .09] |
| Incorrect "Disg" Fear 33% | .06 ± .02, [.02, .09] | .04 ± .02, [.01, .08] |
| Incorrect "Disg" Fear 50% | .09 ± .02, [.05, .12] | .10 ± .03, [.05, .16] |
| Incorrect "Disg" Fear 67% | .09 ± .02, [.04, .14] | .13 ± .03, [.07, .19] |
| Incorrect "Disg" Fear 83% | .10 ± .03, [.05, .16] | .11 ± .03, [.06, .16] |
| Incorrect "Disg" Fear 100% | .08 ± .02, [.03, .13] | .12 ± .03, [.06, .17] |

Performance on the dynamic emotional facial expression task without excluding participants. Values are mean ± standard error of the mean [95% confidence interval]. Disg = disgust.

*Naka-Rushton Modeling*

Figure S4 displays the aggregate Naka-Rushton fits and distributions of parameter estimates. The distribution of the saturation point for fearful faces in the sham condition was largely confined to 1 and thus could not be analyzed. For fearful faces, active sonication increased the slope (*M* = 1.79, *SD* = .61, *CI* = [.79, 3.06], *p* < .001) and offset (*M* = .03, *SD* = .01, *CI* = [.01, .06], *p* = .008) while decreasing the threshold (*M* = .23, *SD* = .13, *CI* = [.01, .50], *p* = .019). For disgust faces, active sonication did not impact the saturation point (*M* = .01, *SD* = .04, *CI* = [-.06, .09], *p* >.250), slope (*M* = .41, *SD* = .58, *CI* = [-.55, 1.59], *p* = .190), threshold (*M* = .01, *SD* = .02, *CI* = [-.03, .05], *p* > .250), or offset (*M* = .02, *SD* =.02, *CI* = [-.02, .07], *p* = .194).

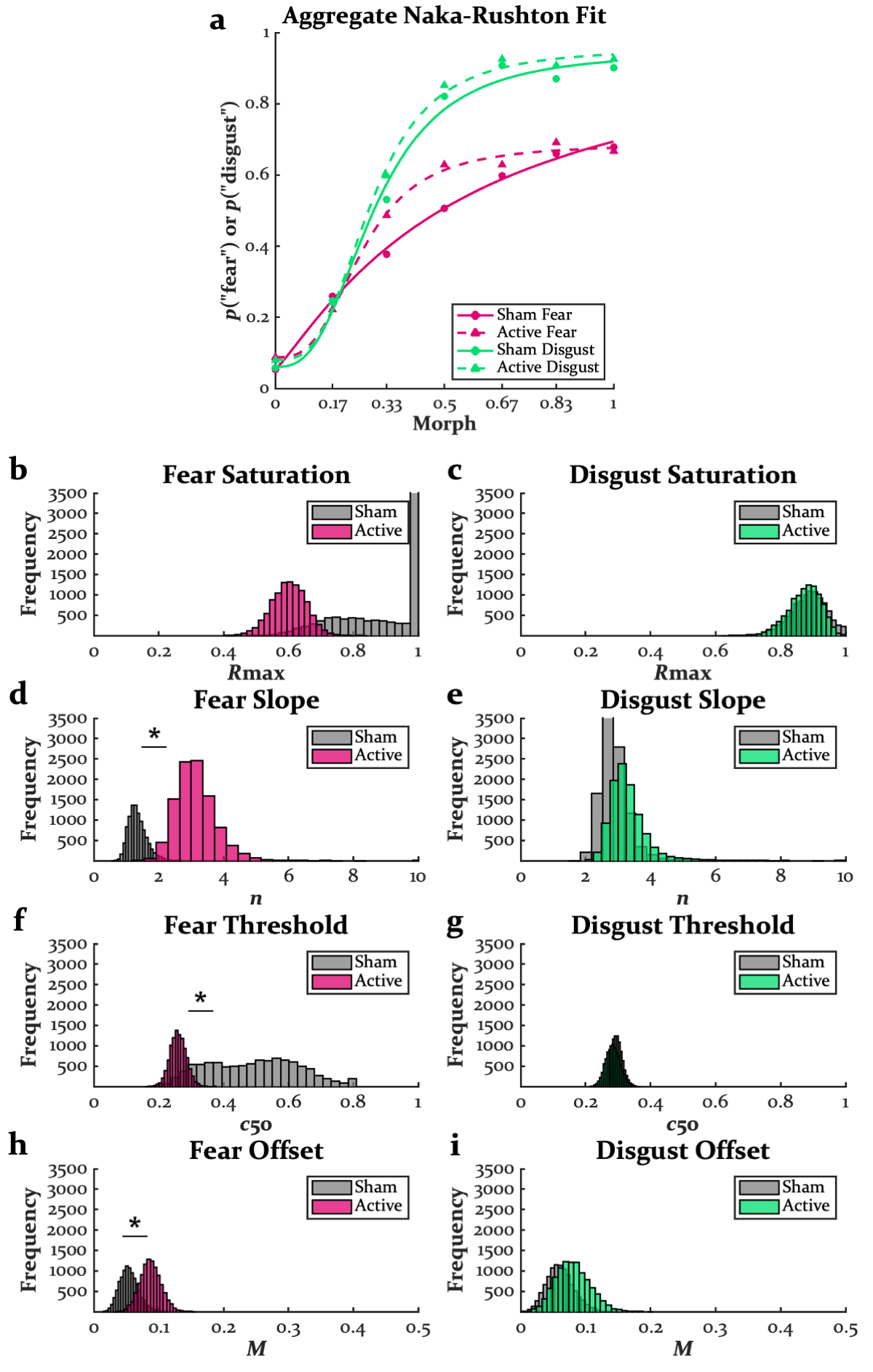

**Figure S4.** Aggregate Naka-Rushton fits (**a**), bootstrap distributions of the saturation point (**b**-**c**), threshold (**d**-**e**), slope (**f**-**g**), and offset (**h**-**i**). * = effect of sonication (*p* < .050).
